# Protein Diffusion Models as Statistical Potentials

**DOI:** 10.64898/2025.12.09.693073

**Authors:** James P. Roney, Chenxi Ou, Sergey Ovchinnikov

**Affiliations:** MIT Computational and Systems Biology; MIT Biology

## Abstract

Machine learning has driven rapid progress in protein structure prediction and design, but key challenges remain such as predicting protein structures when evolutionary information is unavailable, modeling full conformational landscapes, and capturing the thermodynamics of mutations and conformational changes. To address these problems we developed ProteinEBM, an Energy-Based Model of protein conformational space. ProteinEBM’s energies can be used to rank protein structure correctness, predict the energetic effects of mutations, sample from protein conformational landscapes, predict protein structures, and simulate protein folding pathways. Across all of these tasks, ProteinEBM shows performance competitive with or exceeding previous machine learning and physics-based methods, including state-of-the-art performance at predicting the effects of mutations on protein stability.

## 1 Introduction

The use of machine learning has triggered a revolution in protein science and engineering, with AlphaFold often being described as a “solution” to the protein-folding problem [1]. In fact, many aspects of the problem are still open questions. For instance, AlphaFold succeeds at predicting structures for which Multiple Sequence Alignments (MSAs) contain sufficient coevolutionary signal to infer contacts, but protein structure prediction remains difficult when MSAs are shallow or nonexistent. This is a crucial limitation when designing novel proteins, and has constrained designed proteins to much simpler folds than those found in nature [2, 3]. AlphaFold also struggles to accurately predict the structural effects of mutations, since the coevolutionary signal only reflects the wildtype structure. In addition, modern ML methods cannot reliably predict the dynamical pathways by which proteins fold into their native structures. And while important progress has been made toward modeling protein conformational ensembles with machine learning, characterizing the conformational landscapes of proteins with high qualitative and quantitative accuracy, as well as designing proteins with finetuned conformational dynamics, are still open research problems.

A theoretically general solution to these outstanding challenges and others is the development of energy-based models that characterize protein conformational landscapes, as well as efficient samplers for optimizing structures and simulating dynamics from these EBMs. More concretely, energy-based models aim to approximate a function *E*_*θ*_(*x, s*) over protein structures *x* and sequences *s* such that

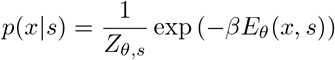

Where *Z*_*θ,s*_ is a generally unknown normalizing constant. Thermodynamically, *E*_*θ*_(*x, s*) is a free energy corresponding to a coarse-grained description of a protein conformational state *x*, integrating out degrees of freedom from the solvent and fine-grained conformational states. There is a rich and highly successful history behind the development of energy functions for proteins, including models based primarily on physical laws, as in the case of molecular dynamics, or a mix of physical terms and knowledge-based terms derived from the PDB, as in the case of potentials like Rosetta [4].

The power and flexibility of EBMs is illustrated by the breadth of their applications. Protein structure prediction corresponds to finding the minimum energy structure *x*^*^ = argmin_*x*_ *E*_*θ*_(*x,s*). Compared with models like AlphaFold, which predicts *x*^*^ in an end-to-end manner, the EBM formulation decouples prediction into the (easier) task of learning a discriminative energy function *E*_*θ*_, followed by the application of an optimization procedure. While the optimization problem is highly nontrivial, the decoupled formulation has the advantage that the compute invested in optimization can be scaled arbitrarily at inference time in proportion to the difficulty of the prediction problem, whereas end-to-end models are fundamentally limited by the scale of the network.

EBMs can also be used to directly simulate coarse-grained protein motion via Langevin dynamics 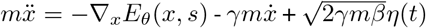. On top of this, EBMs allow for the approximation of free energy differences between conformational states, either by directly calculating energy differences *E*_*θ*_(*x*_1_, *s*)™*E*_*θ*_(*x*_2_, *s*) or by employing enhanced sampling techniques like metadynamics or umbrella sampling. These energy differences encompass important quantities like folding stabilities and binding affinities, as well as the relative occupancies of alternative conformations. The ability to quickly estimate relative occupancies in a differentiable manner also holds great promise in protein design, since it allows sequences to be optimized for the adoption of specific energy landscapes. The breadth of EBMs’ applications also allows them to be trained on diverse sources of data, including protein structures, forces from MD simulations, and experimental measurements of energy differences.

In this paper, we aim to combine the flexibility of EBMs with the expressivity of modern ML architectures. We introduce ProteinEBM, an energy-based model of protein structure and sequence. The architecture of ProteinEBM is inspired by recent advances in protein diffusion and is trained using denoising score matching, but unlike previous protein diffusion models, its learned score function is explicitly computed as the gradient of a learned energy. ProteinEBM’s energy function shows robust performance across a number of tasks, including protein structure ranking, protein stability ranking, protein structure prediction, conformational sampling, and the prediction of folding pathways.

## 2 Preliminaries and Related Work

### 2.1 Molecular Dynamics

Molecular dynamics allows for the simulation of biomolecules with atomic resolution through the forward integration of Newtonian dynamics. The energy functions used to compute atomic forces in MD simulations are called force fields. Force fields are generally composed of simple harmonic and trigonometric terms that define the geometric constraints on bond lengths and angles, as well as electrostatic and dispersive forces mediated by Coulomb’s law and Lennard-Jones potentials. The simplicity of these so-called “molecular mechanics” force fields allows for them to be computed extremely quickly, which is essential since MD simulations must run for billions of timesteps to reach useful timescales. Despite the relative simplicity of molecular mechanics force fields, MD simulations have yielded numerous important insights about the folding and function of biomolecules [5]. Recently, force fields parameterized by neural networks trained on reference quantum mechanical data have enabled simulations of some systems at far greater accuracy, but with even greater computational expense [6].

It is also possible to use machine learning to train coarse-grained models that match the average forces of all-atom simulations on a lower-dimensional representation of a system [7, 8]. This allows for faster simulation, since the coarse-grained energy landscape is significantly smoother. However, existing coarse-grained models trained using force matching require large datasets of MD frames to train, require bespoke prior potentials, and cannot learn from the wealth of experimental and predicted structures in the Protein Data Bank and AlphaFold Database.

### 2.2 Statistical Potentials

In the domain of protein modeling, there is a rich family of models that try to improve the accuracy of molecular mechanics potentials by explicitly incorporating terms derived from the empirical distributions of protein conformations found in the PDB. The most famous method to utilize these “statistical potential” terms is Rosetta, which has been successfully applied to a myriad of tasks in protein modeling and design [4]. In addition to conventional molecular mechanics terms and implicit solvation terms, Rosetta has additional energies based on the Ramachandran distribution for the backbone torsions of each residue, as well as energies derived from the empirical distributions of sidechain rotameric states. Other statistical potentials include Upside, which implements a pairwise distance potential for each combination of two amino acids [9]. Upside was trained using contrastive divergence to minimize the energy of native protein structures relative to alternative samples.

Several papers have also developed creative statistical potentials using modern deep learning methods. Ingraham et al. trained a fully differentiable simulator to learn an energy function that attempts to reproduce experimental structures when optimized under Langevin dynamics [10]. While an impressive engineering feat, this meta-learning approach is computationally intensive and unstable without careful regularization, limiting scalability compared to the far simpler training paradigm of diffusion models. Yang et al. trained an autoregressive model on sequence-structure pairs, and used the model’s structure likelihood as an energy function [11]. While their model has promising structure ranking capabilities, they did not demonstrate *ab initio* protein folding or structure prediction abilities. Du et al. trained a transformer-based EBM to calculate the energies of different sidechain rotameric states and achieved competitive results with Rosetta for rotamer packing accuracy, although their model is not applicable to general backbone conformations [12].

### 2.3 Energy-Parameterized Diffusion Models

Diffusion models have emerged as a leading class of generative models for sampling from complex distributions. These models define a forward noising process that converts samples from the data distribution into an invariant noise distribution:

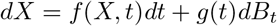

Where *B*_*t*_ is the standard Weiner process and *X*_0_ ∼ *p*_data_ is drawn from the data distribution. By Anderson’s theorem the reverse process is given by

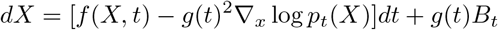

Which converts samples from the noise distribution at *t* = ∞ into samples from *p*_*data*_ at *t* = 0 when integrated backward in time [13]. In a denoising diffusion model, the time-dependent score function ∇_*x*_ log *p*_*t*_(*X*) is approximated by a neural network. Tweedie’s formula states that

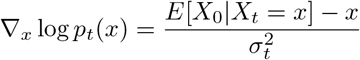

where 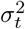 is the conditional variance of the distribution *p*_*t*_(*X|X*_0_ = *x*_0_). This provides a convenient means of training a neural network *s*_*θ*_(*x, t*) to approximate the score function ▽_*x*_ log *p*_*t*_(*x*). While most diffusion models parameterize *s*_*θ*_(*x, t*) as the direct output of a neural network, some authors have noted that is possible to parameterize the score as *s*_*θ*_(*x, t*) = − ▽_*x*_*E*_*θ*_(*x, t*), where *E*_*θ*_ is a learned energy function, which can yield a number of advantages at the expense of increased computational demands from calculating second-order derivatives during training.

Du. et al. used energy-parameterized diffusion models to enable an MCMC-based correction of their diffusion sampler, which allowed them to achieve composability between multiple independent generative models [14]. Thornton et al. used energy-based diffusion models to facilitate composition and low-temperature sampling via MCMC correction, and also introduced various engineering improvements to the energy-training paradigm [15].

Arts. et al. trained an energy-parameterized diffusion model on MD simulations of proteins to act as a coarse-grained forcefield for specific systems [16]. They found the energy-based parameterization to be crucial for obtaining a model that recapitulates atomistic simulation results, since dynamical simulations require forces to be conservative in order to remain stable. Plainer et al. extended this approach with regularization to ensure the reverse diffusion process follows the Fokker-Plank equation, promoting consistency between reverse-diffusion sampling and Langevin dynamics [17]. Jin et al. and Chu et al. both developed energy-based diffusion models for docking pairs of interacting proteins, and showed that the learned energy functions are useful approximations of binding affinity and pose correctness [18, 19]. Concurrently to our work, Sarma et al. used the continuous-time change of variables formula to compute log probabilities from a rigid-docking diffusion model, and showed that they produced energy funnels similar to Rosetta [20]. While the approach is viable for the low-dimensional generative process of rigid docking, integrating the Jacobian trace for log probability computation quickly becomes intractable for higher-dimensional processes.

### 2.4 Conformational Ensemble Generation

Many recent models have aimed to generate samples that reflect the conformational heterogeneity of proteins. Several works have found that subsampling the input MSA to AlphaFold results in diverse structural outputs which can reveal alternative conformations [21, 22]. While highly useful, these alternative structure predictions do not come with any quantitative estimates of relative occupancy, and do not represent unfolded or partially unfolded states. Further, MSA subsampling works by weakening subsets of the evolutionary constraints present in the MSA to produce varied results, which is only effective when the starting coevolutionary signal is sufficiently strong.

Other transferable conformation generators like BioEmu and AlphaFlow have been trained on MD data, allowing them to capture more dynamics than MSA subsampling [23, 24]. These models are very useful tools for MD acceleration, but their lack of an explicit energy function means that there is no direct way of knowing which of their generated samples are more realistic and energetically favorable. Having an energy function built into the diffusion sampler allows us to rerank and resample the structures produced by reverse diffusion, guiding us toward favorable structures that may be sparsely populated by standard diffusion sampling. Compared to standard diffusion models, EBMs can also be used in arbitrary enhanced sampling protocols, like MCMC-corrected sampling and replica exchange between different models, while energy-free methods are limited to enhanced sampling schemes that use additive biasing potentials [25]. Finally, the energies from EBMs can be used to quickly and accurately predict quantities like stability differences, in contrast to models like BioEmu, which must draw thousands of independent samples to estimate free energy differences.

Boltzmann generators use normalizing flows to model conformational ensembles [26]. In theory, Boltzmann generators are the ideal method for ensemble modeling: They provide a normalized probability density (which is strictly more powerful than an energy), and can rapidly produce independent samples. In practice, however, the architectural constraints of normalizing flows have made it difficult to scale Boltzmann generators to large proteins, and most published models only generate small peptides [27]. We see EBMs as a middle ground that combine some of the capabilities of Boltzmann generators with more architectural flexibility.

### 2.5 Structure Scoring Methods

Several machine learning methods have been developed to assess the accuracy of candidate protein structures for a given sequence [28]. Methods like AF2Rank have successfully applied AlphaFold’s confidence scores to identify correct proposal structures for both monomers and protein complexes [29, 30, 31]. While the function learned by AlphaFold to identify correct protein structures may contain some physical principles, it can yield physically unrealistic results when applied to sequence design and mutation effect prediction tasks. Confidence metrics like pLDDT and pTM are also not energies in any mathematically rigorous sense (i.e., they do not approximate the log-density of any relevant distribution), which limits their ability to be used or finetuned for quantitatively accurate dynamics simulation or energy prediction tasks. While log probabilities based on AlphaFold’s distogram predictions can be interpreted as energies, they require AlphaFold to have a largely correct prediction of the native structure in order to be meaningful [32]. AlphaFold-based scoring is also quite slow, especially if many scoring iterations are needed to optimize a structure prediction.

### 2.6 Our Contributions

In this paper, we introduce a powerful diffusion-based EBM for protein modeling. While prior energy-parameterized diffusion models have been applied to specific proteins and rigid docking tasks, we tackle the broader task of learning a universal model of protein conformational energies, with the goal of generalizing to folds and conformations that are not well represented in the PDB. Our model shows robust capabilities across a wide range of tasks, including structure ranking, stability prediction, structure prediction, conformational sampling, and folding pathway prediction. An illustration of our approach can be found in Figure 1.

**Figure 1.**
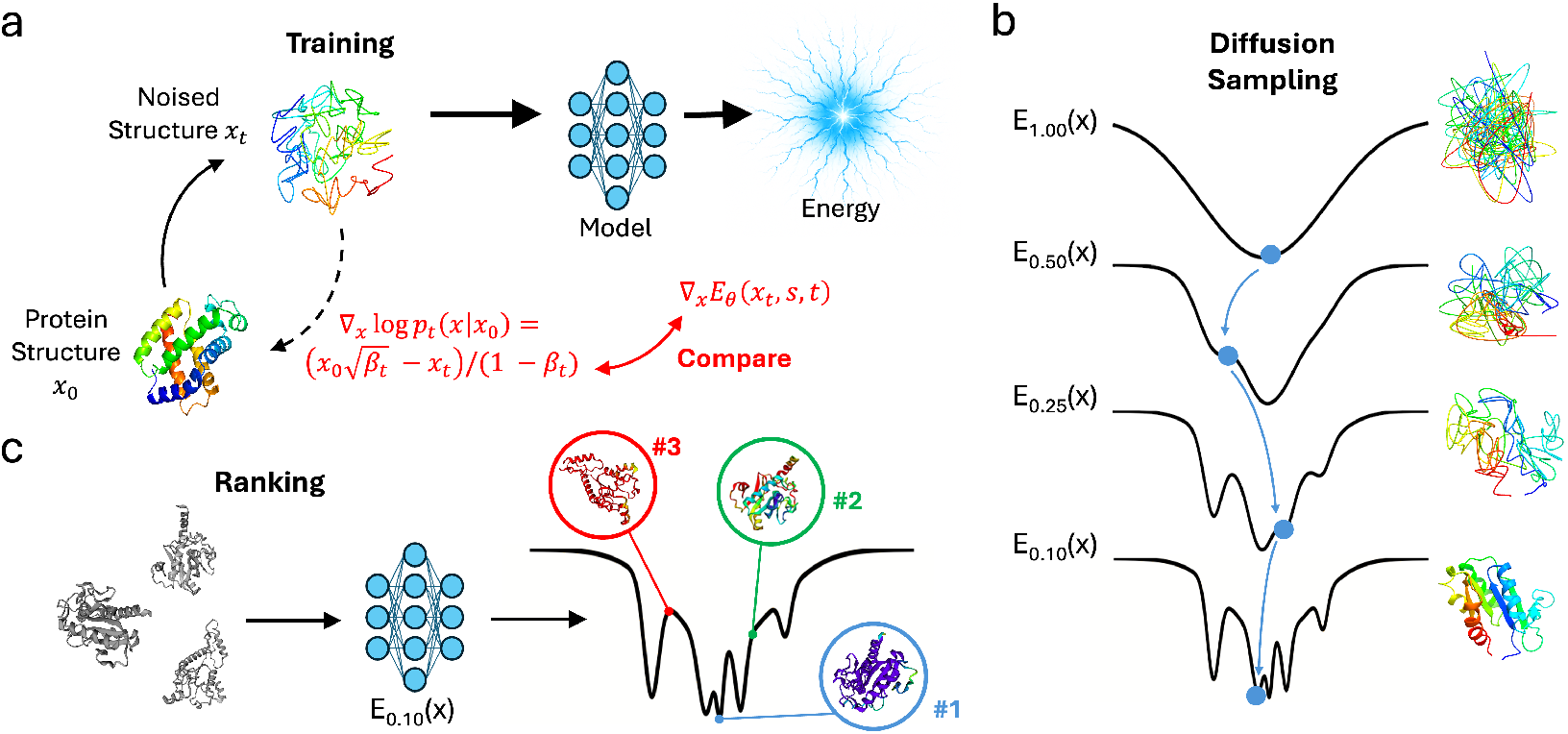
Overview of ProteinEBM. a.) Schematic of the score matching training process. b.) ProteinEBM can generate samples via reverse diffusion over the learned energy landscapes. c.) ProteinEBM can rank arbitrary input structures via its learned energy.

## 3 Methods

ProteinEBM is an energy-parameterized, sequence-conditioned protein diffusion model based on the diffusion modules from AlphaFold3 and Boltz-1 [33, 34]. This means that, unlike previous diffusion models, ProteinEBM does not utilize an inherently equivariant model architecture, and relies on data augmentations to learn 3D symmetries. We found a non-equivariant architecture to be superior to Invariant Point Attention for achieving stable denoising trajectories with an energy-parameterized model, likely because optimizing second-order derivatives through IPA can result in instability. We downscaled the model from the version used in Boltz-1, resulting in a model with 85M parameters.

We trained ProteinEBM on a set of protein domains from the PDB and AlphaFold Database, using domain definitions from CATH and The Encyclopedia of Domains (TED) [35, 36]. We pretrained on a total of 32k CATH domains, 590k AFDB domains derived from the dataset of Lin et al., as well as 18k two-chain protein complexes from the set published by Jin et al. [18, 37]. Following pretraining, we finetuned a subset of our models on molecular dynamics simulations of 1k CATH domains that were previously used to train BioEmu [23]. These simulations were run at 300K using various AMBER forcefields, and contain both folded and unfolded structures.

We pretrained our models for at least 60 epochs, with each pretraining epoch containing all 32k CATH structures, all 18k complex structures, and 32k AFDB structures. We then finetuned on up to 3M frames of MD data, with the exact amount of finetuning varying depending on the application. We found that MD finetuning was unnecessary for obtaining strong performance on structure ranking and stability prediction tasks, but did help increase the diversity of generated samples.

ProteinEBM was trained using the denoising score matching framework. For a sample *x*_0_ ~ *p*_*data*_ and its noised version 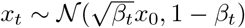, the score matching loss is

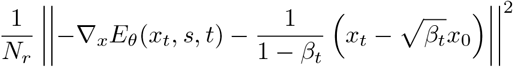

Where *N*_*r*_ is the number of residues. Optimizing this loss drives *E*_*θ*_(*x*_*t*_, *s, t*) to equal − log *p*_*t*_(*x*| *s*) up to a constant, where *p*_*t*_(*x s*) is the sequence-conditional data density noised to level *t*. At low enough *t*, the learned energy should theoretically approach − log *p*_*data*_(*x*| *s*). However, this ideal outcome is not guaranteed, and whether this denoising task results in an energy that usefully resembles the true free energy is ultimately an empirical question.

As mentioned previously, AlphaFold-based scoring can suffer from pathological modes in which proteins with exposed hydrophobic residues or non-compact structures are scored confidently, despite their physical unfavorability. This is likely due to AlphaFold implicitly inferring the presence of an unseen binding partner or domain, since the model was trained on cropped portions of single chains with many interactions missing. To avoid this pathology in ProteinEBM, all training residues with a cropped out interaction partner were marked with an “external contact flag” that was passed into the model as input. At inference time these flags are set to zero, keeping the model from inferring any extra context.

## 4 Results

### 4.1 Decoy Ranking

Discriminating correct native structures from plausible but incorrect “decoy” structures is a fundamental capability for energy-based protein models. This capability is important for selecting models generated by structure predictors and weighting conformational ensembles produced by MSA subsampling methods, as well as for predicting protein structures in regimes where coevolutionary information is scarce. In this setting, where tools like AlphaFold can fail, investing a larger amount of compute searching over structure space for low-energy structures is a promising approach to improving prediction results [29]. This is similar to the idea of test-time scaling in Large Language Models, for which optimization over EBMs has recently been proposed as a general solution [38].

To validate ProteinEBM’s ranking ability we used the Rosetta decoy set, which contains 133 native protein structures with thousands of decoys for each native structure [39]. We subdivided the dataset into 50% validation and 50% test structures, and filtered our training data to have less than 40% sequence identity to proteins in either set (normalized by the length of the shorter protein to prevent the inclusion of subsequences). Additionally, we performed a structure-based split against 20 test targets by filtering out all training structures sharing a topology-level CATH classification with the targets. We refer to the proteins subjected to the CATH split as hard targets, and those subjected to the sequence split as easy targets. For our initial attempt at decoy ranking, we used a ProteinEBM model finetuned on 1M MD frames with only the first and last transformer layers unfrozen. To rank a decoy structure *x* we computed *E*_*θ*_(*x, s, t*), with the time level *t* becoming a hyperparameter.

With this protocol, we found that decoy ranking performance peaked at time levels slightly above 0, and then decayed for larger values of *t*. Setting *t* = 0 should theoretically make the model energy converge to the data density, but in practice at extremely low noise levels the model only needs to correct very local structural features and thus its energy does not correlate well with global structural correctness. At very high levels of *t* the model energy converges to a Gaussian density, so it effectively ranks decoys based on their radius of gyration. Low but nonzero values of *t* interpolate between these extremes and give the best performance. After our initial training, we found that decoy ranking performance on the validation set peaked at times between t=0 and t=0.15 (Figure 1b). Based on this observation we then trained an “expert model,” referred to as ProteinEBM-x, only on times below t=0.15, which significantly improved performance by concentrating model capacity on the relevant noise levels. The exact training protocols for our models can be found in Appendix A. Unless otherwise stated, we use ProteinEBM-x in all reported results with t=0.05 as the time level, which we chose to optimize ranking ability on the validation set.

ProteinEBM-x performed very well on ranking decoys in the test set, with an average Spearman correlation of the 0.838 between the model energy and decoy TMScore [40]. This is significantly higher than the Rosetta energy function, which had an average Spearman of 0.757 (*p* = .0005). The average TMScore of ProteinEBM-x’s minimum-energy decoy for each target was 0.905, compared to Rosetta’s top-1 TMScore of 0.899 (*p* = 0.46, n.s). Our models consider the sequence-structure relationship and do not merely use backbone realism for ranking, as performance degrades significantly when the sequence is masked out (Figure 2b). Energy-based ranking also far outperforms a simple strategy of ranking structures by the norm of the score, highlighting the necessity of the energy parameterization (Figure S1). We found no significant performance degradation on targets subjected to the stricter CATH-based split (Figure 2c-d), indicating good generalization.

**Figure 2.**
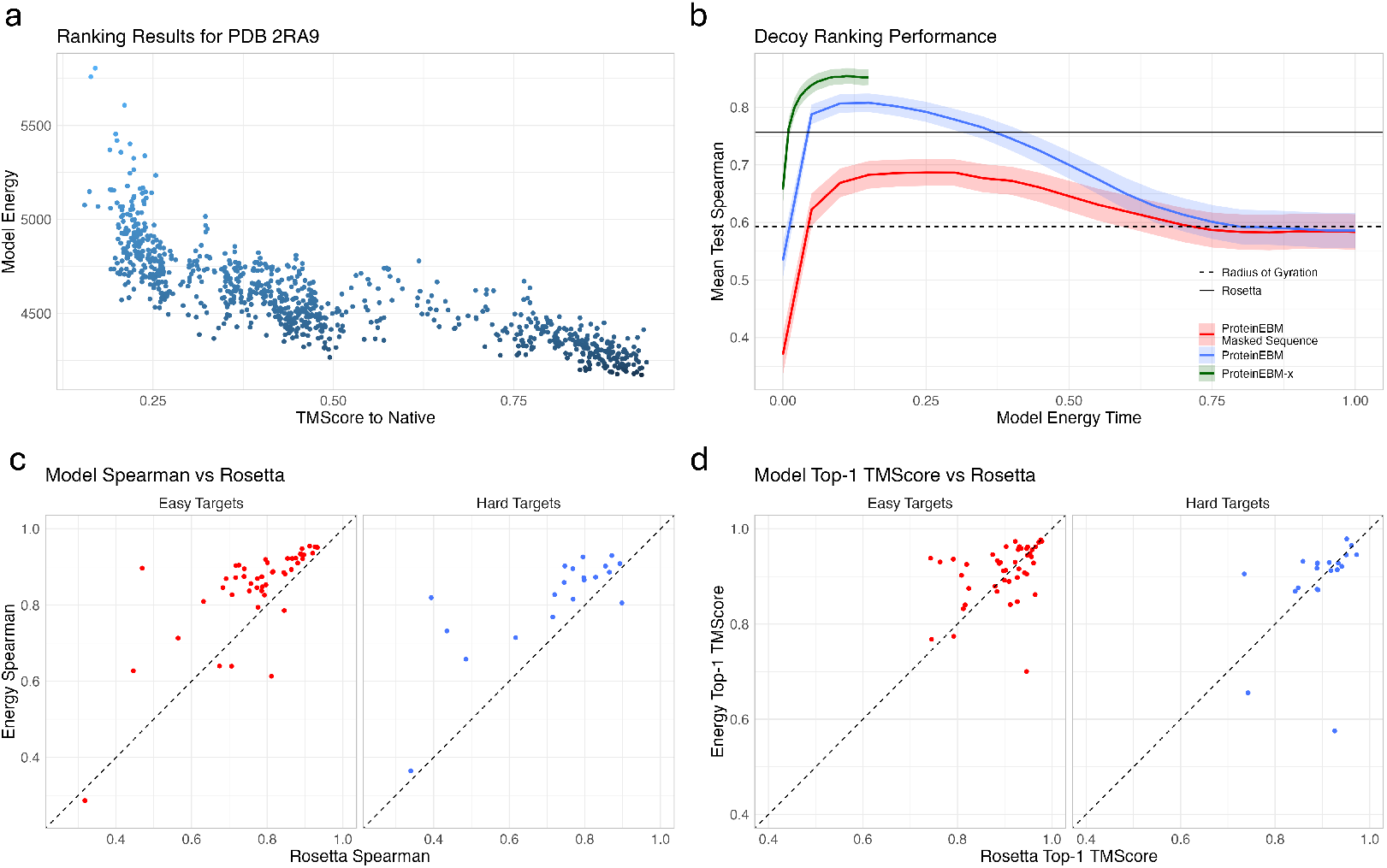
ProteinEBM-x can be used to accurately rank protein structure accuracy. a.) Example correlation between ProteinEBM-x energy and decoy TMScore for a test set protein. b.) Mean Spearman correlations between energy and TMScore at different values of *t*. Ribbons are standard error of the mean. c.) Comparison of Spearman correlations for ProteinEBM-x and Rosetta on each target in the test set. Dots are separated into easy and hard targets. Hard targets were subjected to a structure-based split and are farther from the training set d.) Comparison of the TMScore of the top-ranked decoy for each test target for ProteinEBM-x and Rosetta.

### 4.2 Stability Prediction

We next examined the ability of ProteinEBM-x to predict stability changes in proteins upon mutation. We did this by evaluating the energy difference

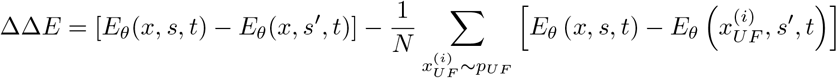

Where the right represents a heuristic Monte Carlo estimate of the free energy difference integrated over the unfolded states. We implemented *p*_*UF*_ as a Ramachandran-random sample of backbone torsion angles. This is only an approximation, and requires that the folded and unfolded conformational ensembles remain nearly constant under mutation. While not fully rigorous, such an approximation is commonly used when estimating point mutation energies [18], [7].

We benchmarked our predicted ΔΔ*G*s on the stability datasets in ProteinGym, most of which were computed by Tsuboyama et al. via cDNA proteolysis [41, 42]. Our estimated energy differences showed a very robust correlation with experimental values. With an average Spearman correlation of .684, ProteinEBM-x outperforms all previously benchmarked models and sets a new state-of-the-art on zero-shot stability ranking on ProteinGym (Table 1). The previously highest-performing models are all structure-aware protein language models like ESM3, which has over 15 times as many parameters as ProteinEBM-x [43]. In contrast, ProteinEBM-x is an energy-based structure denoiser that was never explicitly trained to predict masked amino acids, and thus represents a new paradigm for accurate, parameter-efficient unsupervised stability prediction. ProteinEBM’s success at this task also stands in contrast to confidence metrics from AlphaFold-like structure predictors, which generally exhibit mediocre correlation with stability changes.

**Table 1:**
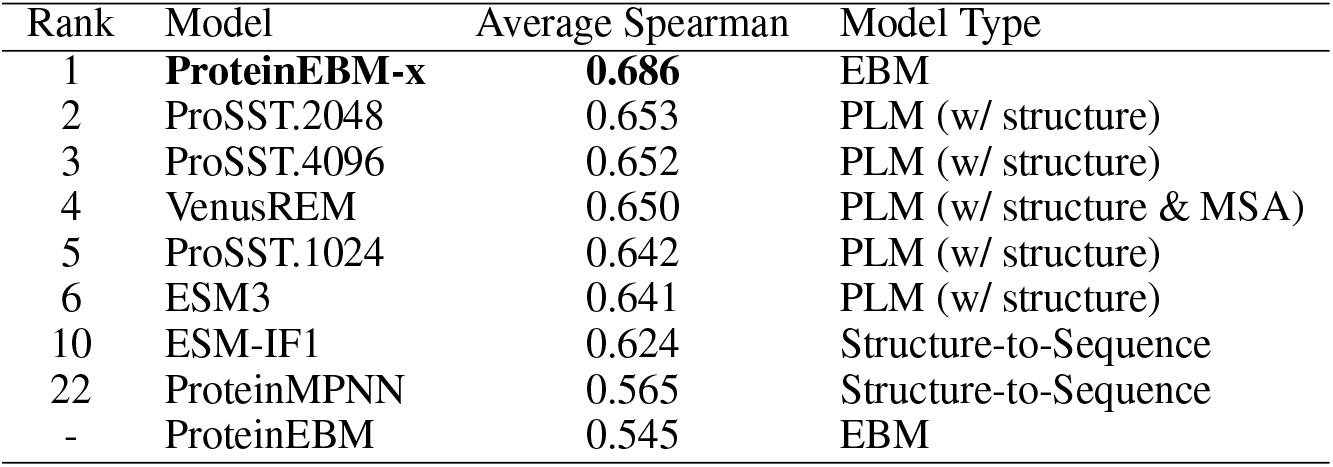
State-of-the-art performance on the ProteinGym stability benchmark.

To expand the number of proteins used for evaluation we supplemented ProteinGym with the remaining datasets from Tsyuboyama et al. that were not included in the original benchmark, many of which are *de novo* proteins [41]. Adding these extra proteins resulted in an even larger performance gap between ProteinEBM-x and ESM3, which we used as a baseline on the expanded set. This was likely due the addition of more *de novo* proteins with no evolutionary history, as the performance gap between ProteinEBM-x and ESM3 was correlated with MSA depth (Figure 3d). For proteins with 0 or 1 MMSeqs2 hits, which likely all represent *de novo* proteins, ProteinEBM-x improved on ESM3 by an average of .178 Spearman points (.697 vs .519, *p <* 0.0002). This suggests that ESM3’s performance is more dependent on evolutionary conservation relative to ProteinEBM-x.

**Figure 3.**
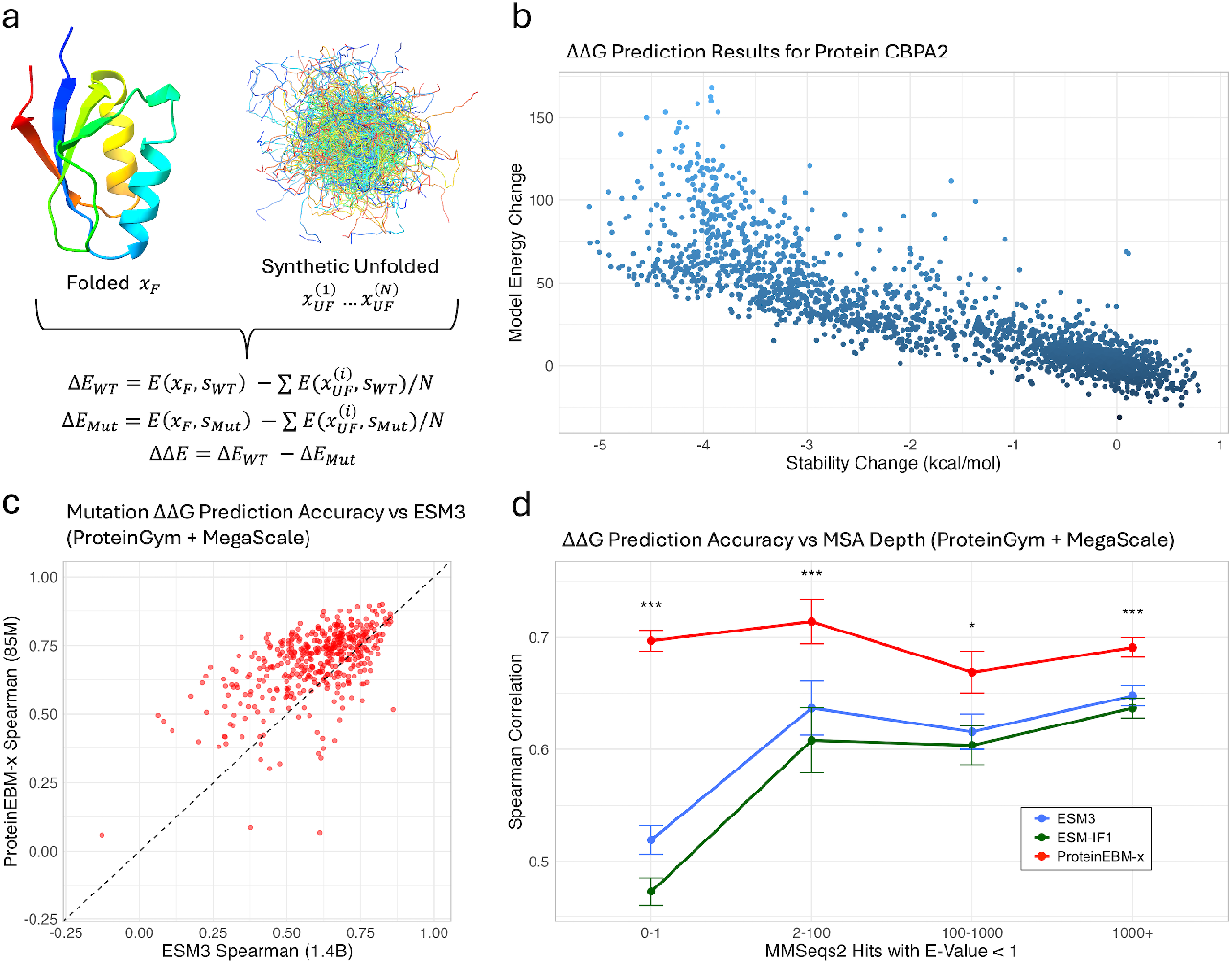
ProteinEBM-x energies accurately predict changes in protein stability. a.) Method for approximating ΔΔ*G* with folded and unfolded structures b.) Example correlation between predicted and experimental ΔΔ*G* for protein CBPA2. c.) Spearman correlations between ProteinEBM-x energy differences and experimental ΔΔ*G*s vs ESM3 Spearmans for all stability targets in ProteinGym and MegaScale c.) ProteinEBM-x, ESM3, and ESM-IF1 Spearmans on ProteinGym and MegaScale v.s. the MSA depth of each protein. Error bars are standard error of the mean.

### 4.3 Conformational Sampling

Our model can be used to generate structural samples in two ways: either by performing i.i.d. sampling via the reverse diffusion process, or by exploring the local energy landscape via dynamical simulation. The former method allows for perfectly decorrelated samples to be drawn without concern for the convergence of the sampling process, while the latter allows for local exploration of specific regions of conformation space and the elucidation of transition paths and kinetic parameters. Simulating directly from a low-noise energy level may also reveal new energetic basins that have been learned by the low-noise model, but are never revealed in reverse diffusion due to the divergence of the higher-noise energies from the Fokker-Planck equation. We performed local sampling using Langevin dynamics, and the details of our simulation algorithm can be found in Appendix B.1.

To examine ProteinEBM’s ability to predict structures and sample conformational landscapes, we examined its performance on 11 fast-folding proteins first examined by Lindorff-Larson et al. [5]. These proteins were also split from the training set at a sequence identity threshold of 40%.

To generate structural landscapes, we performed Langevin annealing trajectories, in which we started from random noise and took 100 Langevin steps at each noise level before taking a reverse diffusion step to the next noise level in the reverse diffusion process. This approach enables better exploration by allowing the samples to equilibrate to each energy level. The ProteinEBM base model was used for timesteps with *t >* 0.1, with ProteinEBM-x used below *t* = 0.1. To analyze our models’ exploration of conformational space, we used Time-Lagged Independent Components Analysis to compare our samples to the simulations published by Majewski et al [8].

We found that, with Langevin annealing, our model generally captured the correct native structure. For all but 1 target the lowest-energy samples were within 3.5 Angstroms RMSD of the native structure. The only exception was the WWDomain, for which the lowest-energy structure exhibited all of the correct secondary structure arrangements but had some differences in loop conformations. In addition, NuG2 also shows a deep energy well at a strand-swapped nonnative conformation, which appears as the lowest-energy sample in some annealing runs (Figure S2). For all targets, the minimum-RMSD structure sampled during annealing was within 2.5Å of the native structure. Interestingly, we found that the sampled structures often had significantly lower energies than the native structures themselves, but this difference disappeared after subjecting the native structure to a fast relaxation protocol with the ProteinEBM-x energy.

For some targets the lowest-energy samples were highly accurate despite the fact that most samples were unfolded or otherwise distant from the native. For example, the vast majority of sampled TrpCage structures were unfolded, but the lowest-energy sample was 2.54Å from the folded NMR structure. This highlights the power of EBMs to perform massive sampling and subsequent reranking to facilitate accurate structure prediction, which is not possible with models like BioEmu that lack an explicit energy. The energy funnels in Figure 4 are also qualitatively reminiscent of energy funnels produced by methods like Rosetta.

**Figure 4.**
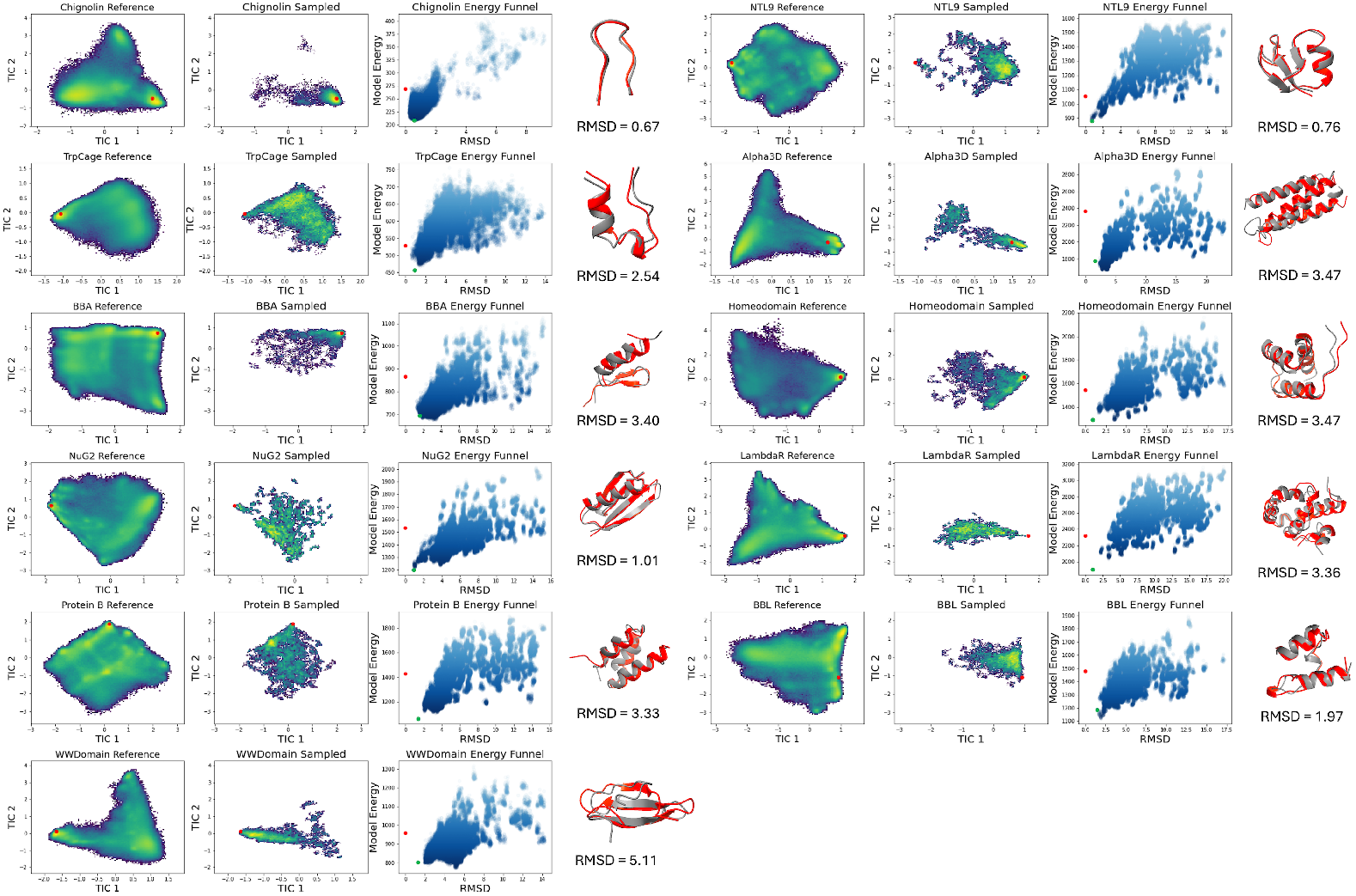
ProteinEBM effectively samples conformations of fast-folding proteins with Langevin annealing. TICA plots showing conformational landscape for MD simulations (left) and ProteinEBM samples (right) for each fast folder, with red dot indicating native structure. Energy funnels show the relationship between model energy (*t* = .05) and RMSD to native for sampled structures. For funnel plots, the red dot indicates the native structure and green dot indicates the native after running a fast relaxation procedure. For protein structure graphics, the lowest-energy ProteinEBM sample is in red, and the native is in gray.

In addition to capturing the native structures our model explores a significant portion of the non-native conformational space, although not as much as all-atom simulation. This not unexpected because ProteinEBM is a native-centric model with limited exposure to unfolded states in its training data, and its MD finetuning data were simulated at 300K while the reference simulations were run at 350K.

### 4.4 Folding Simulations

While Langevin annealing simulations allow for some exploration of conformational space, they cannot capture the pathways by which proteins fold, since the samples are generated by transforming from a Gaussian noise distribution rather than the distribution of the unfolded ensemble. However, ProteinEBM and ProteinEBM-x are also capable of performing direct folding simulations by running Langevin dynamics at a low noise level starting from an unfolded state. To investigate this capability, we performed Langevin simulations starting from Ramachandran-random chains using the energy function from ProteinEBM-x at *t* = .05. The simulations thus take place on a manifold of protein structures with a small amount of Gaussian noise, but the noise can be removed to produce a final trajectory by projecting each noisy simulation step to *t* = 0 using the model’s predicted score.

To assess the accuracy of our folding simulations, we folded Protein G, its engineered variant NuG2, and Protein L, all of which have extensively characterized folding pathways. These bacterial immunoglobulin-binding proteins all have the same topology consisting of a single helix packed against N- and C-terminal hairpins. Experimental techniques like *ϕ*-value analysis have shown that the C-terminal hairpin adopts a more stable, native-like fold at the transition state in Protein G, while Protein L primarily folds through the N-terminal hairpin [44]. NuG2 was engineered starting from Protein G, and the N-terminal hairpin was stabilized to make its folding pathway resemble that of Protein L [45].

For each protein, we ran a sufficient number of simulations to observe >20 independent trajectories that folded to the native state within 3Å RMSD. We ran 1200 simulation trajectories for Protein G, 800 for NuG2, and 4800 for Protein L. For Protein G and NuG2 the lowest-energy sampled structures had the native fold, while the lowest-energy structure for Protein L was a strand-swapped version of the native structure (Figure S3). To analyze the folding trajectories we extracted transition intervals in which the simulated chain went from >10 Å from the native structure to <3 Å from the native or vice-versa. To summarize the sequence of native contact formation during folding and unfolding we computed the fraction of time native contacts were present along transition paths. A higher value indicates that a contact tends to form earlier during folding and break later during unfolding.This analysis was inspired by Lindorff-Larsen et al. and Chang and Perez [5, 46].

We found that, consistent with experiment, the C-terminal hairpin contacts had the greatest presence along transition paths in Protein G, while NuG2 and Protein L exhibited folding pathways that were shifted toward the formation of N-terminal contacts (Figure 5). While it is difficult to make quantitative inferences about folding pathways from our simulations (which may have physically unrealistic kinetics due to coarse-graining), the qualitative agreement with experiments highlights the potential of EBM-based simulation for exploring the process of protein folding.

**Figure 5.**
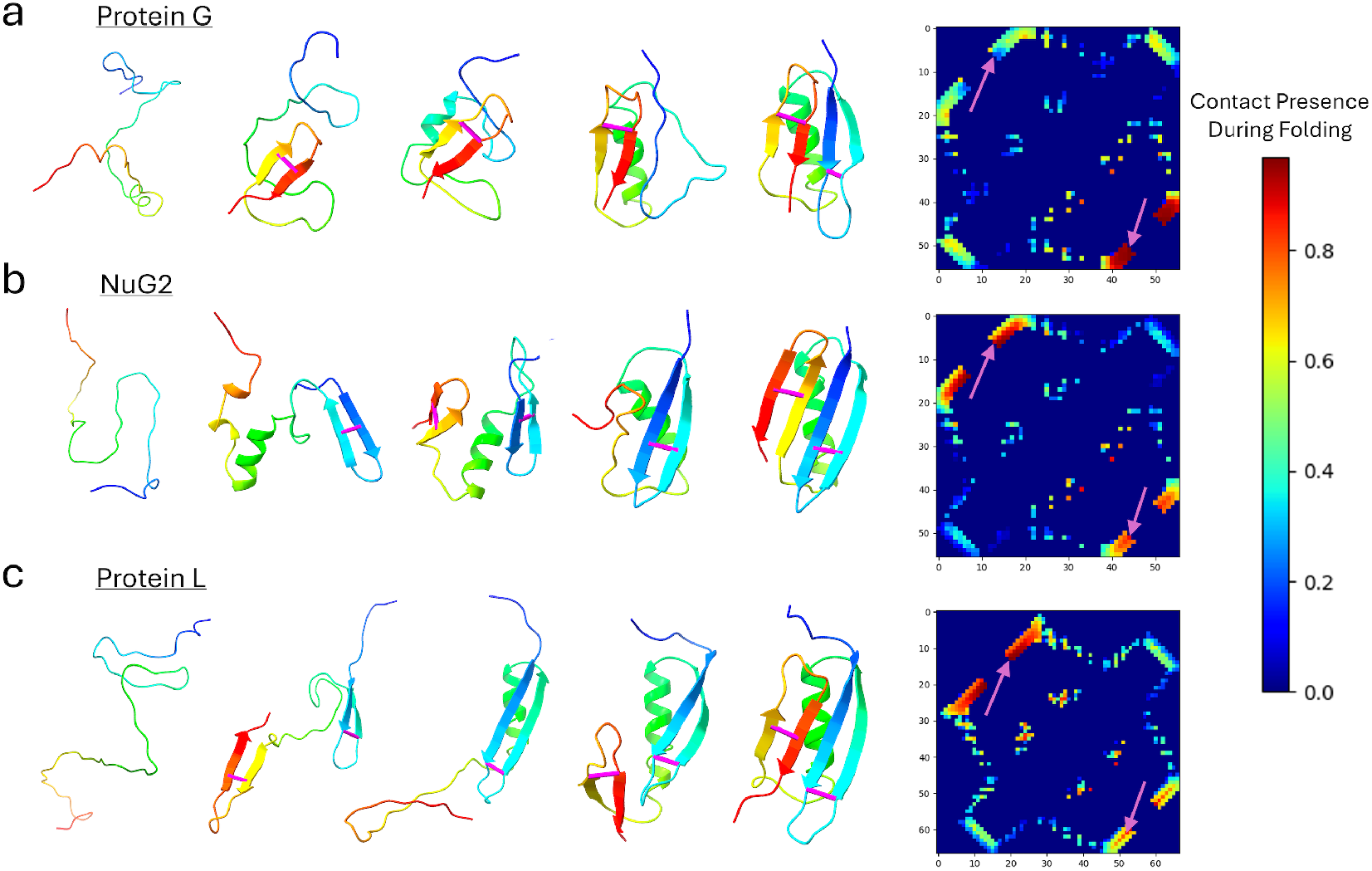
ProteinEBM-x can be used to rapidly simulate folding trajectories of proteins. a.) Left: An example folding trajectory for Protein G, with snapshots taken at different frames from the simulation. Formation of example contacts labeled in Magenta. Right: Contact presence fraction during folding transitions, with example contacts indicated with magenta arrows. b.) Same as part a for NuG2. c.) Same as part a for Protein L.

### 4.5 Structure Prediction

Finally, we tested ProteinEBM’s ability to predict structures for proteins in our Rosetta test set without using MSAs. In our initial experiments on the validation set, we found that using models finetuned on more MD data facilitated better exploration. Ultimately, we settled on a protocol of running Langevin annealing with a ProteinEBM model finetuned on 3M frames of MD data, followed by resampling the lowest-energy structures and re-annealing with the a ProteinEBM-x model that was not trained on any molecular dynamics data, which was also used to score the final outputs. As an optional step to improve ranking and increase backbone quality, we tried clustering the final structures and passing the lowest-energy cluster centers to AF2Rank. Despite the high computational cost of AF2Rank, this step was computationally cheap because we scored at most 40 samples per target using AF2Rank out of thousands of total ProteinEBM samples. We selected the final prediction to be the AF2Rank output structure with the highest pTM. Details of the sampling protocol can be found in Appendix B.2.

To create strong baseline methods, we ran AlphaFold2 in single-sequence mode for 10,000 seeds per target, and AlphaFold3 in single-sequence mode for 15,000 seeds per target, taking the prediction with the highest pTM as the final output. We calculated these sample numbers based on empirical measurements of the runtime for each method. Details of the calculation are provided in Appendix H.

On the easy targets (split from the ProteinEBM training data at 40% identity), ProteinEBM sampling outperformed both the AF2 and AF3 baselines (Figure 6c-d). The lowest-energy samples from ProteinEBM had an average TMScore of .613, compared to top-1 TMScores of .584 for AF2 and .529 for AF3 (*p* = .43 and .01, respectively). AF2Rank refinement on the low-energy ProteinEBM samples further boosted the average top-1 TMScore to .690 (*p* = .001 and *p <* 0.0002 vs AF2 and AF3). While these targets are “easy” relative to the other test targets split by topology, they are still absent from the ProteinEBM training set and present in the training sets of AF2 and AF3, underscoring the effectiveness of ProteinEBM at large-scale sampling and ranking.

**Figure 6.**
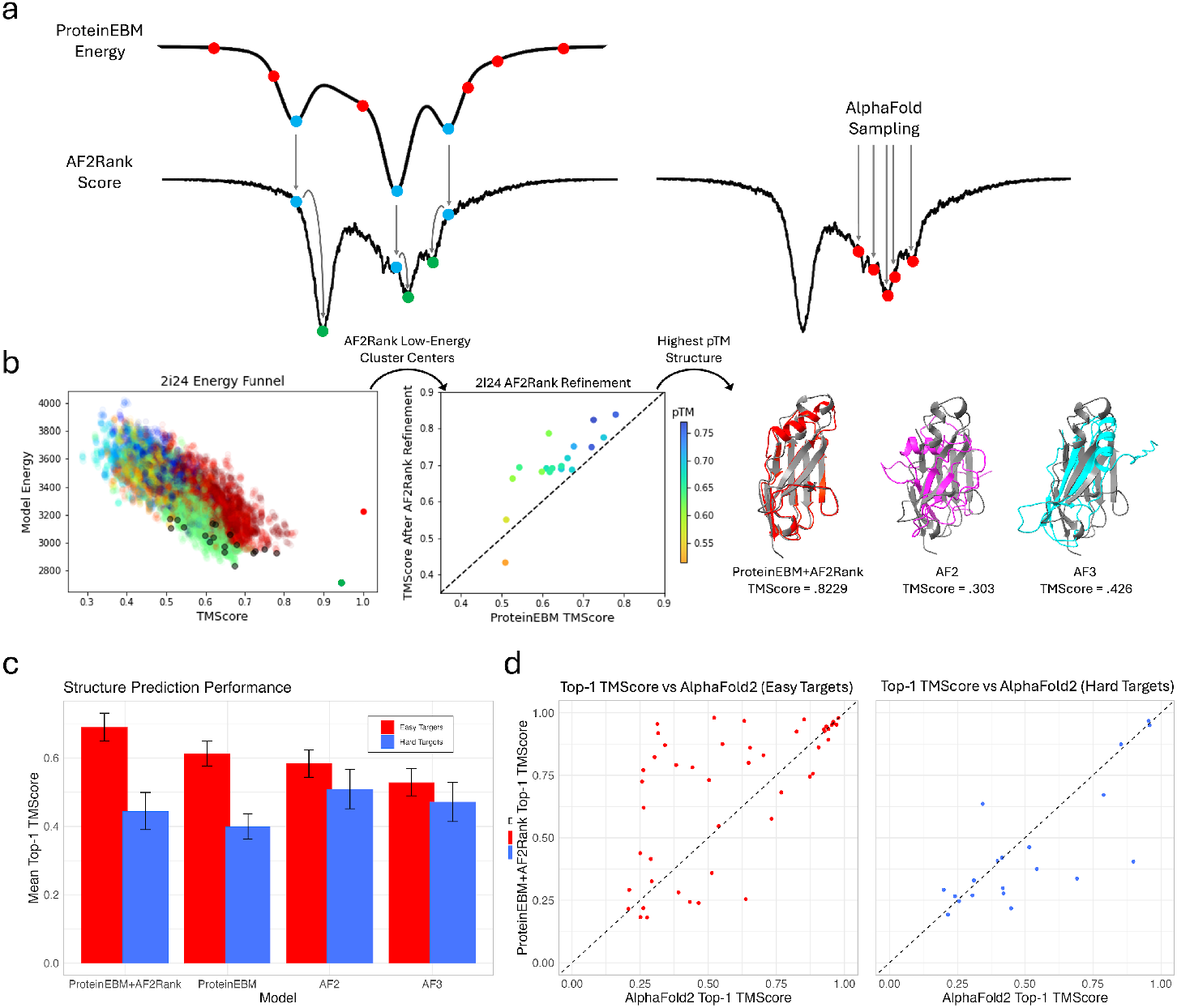
ProteinEBM and ProteinEBM-x can be used to sample and score structure predictions. a.) ProteinEBM Langevin annealing (left) explores broadly, and the best samples are passed to AF2Rank. Standard AlphaFold sampling (right) can fail to sample global minima. b.) Left: TMScore vs Energy for ProteinEBM samples on PDB 2i24. Red dot is native, green dot is relaxed native, colors are clusters, gray dots are cluster centers. Center: AF2Rank refines and reranks lowest-energy cluster centers. Right: Predicted vs native structure for ProteinEBM+AF2Rank, AF2, and AF3. c.) Average top-1 TMScore for ProteinEBM, ProteinEBM+AF2Rank, AF2 and AF3, with performance split between easy and hard targets. Error bars are standard error of the mean. d.) Scatter plot of top-1 TMScores for ProteinEBM+AF2Rank vs AF2, split by easy and hard targets.

On the hard targets split from the ProteinEBM training data by topology, ProteinEBM sampling underperformed AF2 and AF3 regardless of AF2Rank refinement. Since decoy ranking performance remained strong on these targets (Figure 1c-d), this performance degradation suggests that Pro-teinEBM is capable of ranking the accuracy of structures far outside the training distribution, but has a harder time sampling unknown folds.

On unsuccessful prediction targets ProteinEBM exhibited multiple failure modes. Sometimes the model failed to explore structures close to the native, despite the fact that the native structure was scored with a lower energy than the structures produced by the model. In other cases, the model explored native-like structures but erroneously ranked other worse structures with lower energy. Examples of these failure cases can be found in Figure S9. These failure modes point to improvements in both the model’s ranking ability and exploration ability as important areas of future work. Broader exploration could also benefit from the use of generative models that explicitly enumerate diverse protein folds.

## 5 Discussion and Future Directions

We have developed an energy function for protein structural landscapes that shows strong performance in structure ranking, protein structure prediction, and conformational sampling, as well as state-of-the-art performance in zero-shot protein stability ranking. Our EBM is trained using a simple denoising score-matching paradigm, and is highly efficient compared to frontier structure prediction models. Our strong results demonstrate that EBMs are a promising framework for grounding ML protein structure models in thermodynamic principles. Protein conformational preferences, stabilities, binding affinities, and coarse-grained dynamical motions can all be formulated in terms of free energies, which allows EBMs like ours to generalize to a wide array of tasks in protein modeling. And by decoupling structure scoring from sampling, we can scale computing resources arbitrarily at inference time to facilitate a broader search over structures for difficult prediction targets. This paradigm can potentially relax the dependence of structure prediction on multiple sequence alignments, thereby unlocking greater regions of protein fold space for *de novo* design.

There are many possible avenues for improving the capabilities of ProteinEBM. While ProteinEBM-x exhibits state-of-the-art unsupervised performance on ΔΔ*G* prediction, large experimental stability datasets could be used to actively supervise the model’s energy differences. The model could also be finetuned using contrastive divergence, with large sets of decoys generated from sampling serving as a replay buffer of contrastive samples. Our models’ capabilities could be evaluated and optimized for protein complexes. Future work should also explore the use of more enhanced sampling techniques to improve exploration during structure prediction and simulation. It will also be useful to train EBMs on both finer and coarser levels of protein structure representation, such as all-atom models and models that operate on smoother latent representation spaces.

## 6 Code and Data Availability

All code and model weights are available at https://github.com/jproney/ProteinEBM.

## 7 Acknowledgements

James P. Roney acknowledges support from the Fannie and John Hertz Foundation. Chenxi Ou acknowledges support from the Herman N. Eisen Fellowship Fund. We thank all members of the Ovchinnikov lab for useful discussions and feedback. Sergey Ovchinnikov acknowledges funding from NSF grant MCB2032259, CocaCola and Amgen.

## A Training Details

The primary score-matching loss used for training ProteinEBM is as described in Section 3:

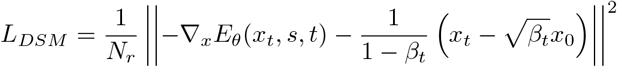

We parameterized the energy output as *E*_*θ*_(*x*_*t*_, *s, t*) = Σ_*i*_||*r*^(*i*)^(*x*_*t*_,*s,t*)||^2^,where *r*^(*i*)^(*x*_*t*_, *s, t*) is a 3-dimensional output vector corresponding to each residue.

As an auxiliary prediction, we also added a non-conservative direct score output to the model. The auxiliary score loss was computed as:

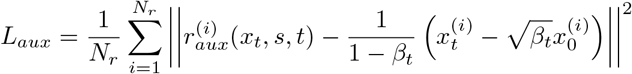

Both the *r* and *r*_*aux*_ projections were implemented as linear projections from the final transformer layer. Finally, we introduced an all-atom modeling loss as follows:

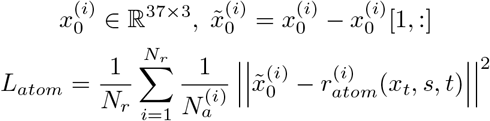

Where *N*_*a*_ is the number of non-*Cα* heavy atoms present in the residue. Effectively, 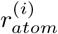 aims to predict the *Cα*-centered coordinates of all heavy atoms in the residue from the noised *Cα* coordinates *x*_*t*_. Note that, because the non-*Cα* atoms are not part of the energy function or the diffusion process, predictions from the all-atom output head suffer from mean collapse and do not necessarily correspond closely to the data distribution. This can potentially be corrected by including sidechain atoms in the diffusion process or adding losses that penalize physical violations. The final training loss was computed as

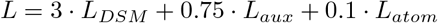

We trained using the Adam optimizer with base learning rate .0002 and a OneCycle learning rate schedule. During training the sequence information *s* is masked out 10% of the time. We also trained with self-conditioning 50% of the time as introduced by [47]. When self-conditioning was used, we obtained a denoised estimate 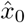 using the auxiliary score output from a preliminary model run, and then recycled it back into the model input for the main training step. During decoy scoring we passed the candidate structure coordinates into both the main input channel and the self-conditioning channel.

The hyperparameters for our cartesian diffusion process are largely derived from Yim et al. [48]. The marginal distribution of our diffusion process at time *t* is given by 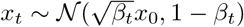 where

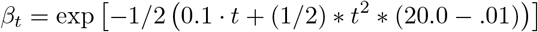

The diffusion process and score are all defined in units of nanometers in order to make the data samples closer to unit variance.

Our base ProteinEBM models were pretrained for 60 epochs with *t* sampled uniformly between 0 and 1. ProteinEBM-x was pretrained for 60 epochs with *t* sampled uniformly between 0 and 0.15, followed by 20 epochs of finetuning with *t* ∼ min(𝒩(.05, .025), 0.15), and further finetuned on 1M MD frames (unfreezing only the first and last layers) with the same time distribution.

## B Sampling and Simulation

### B.1 Overdamped Langevin Dynamics

For the simulation of local dynamics, we utilized overdamped Langevin dynamics on the energy surface for time-level *t* = *τ* defined as follows:

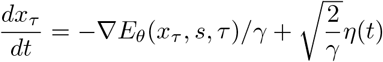

Where *γ* is a friction coefficient and *η* is decorrelated Gaussian noise. When discretized as an Euler-Maruyama step, this becomes:

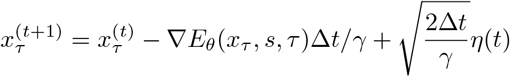

Now compare this with one step of denoising diffusion at time *τ*, followed by re-noising back to time τ

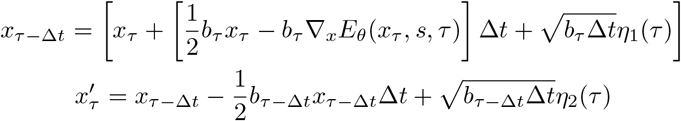

For a standard variance-preserving forward diffusion process with drift coefficient 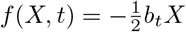 and diffusion coefficient 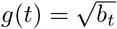. In the limit of small Δ*t* this converges to

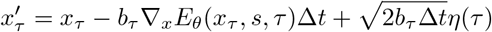

Which is equivalent to overdamped Langevin dynamics with friction coefficient *γ* = 1*/b*_*τ*_. Thus, we can sample from the energy surface at time *τ* by iteratively running reverse and forward diffusion steps with a small timestep Δ*t*. The energy in the reverse diffusion step can also be adjusted with a correction factor *T*_*train*_*/T*_*sim*_ that approximately shifts the simulation temperature from the training temperature, although this is not fully rigorous since the free energy itself is temperature-dependent. The correspondence between Langevin dynamics and iterative partial diffusion is analogous to the result from Arts et al. translated into the language of continuous-time diffusion [16].

### B.2 Sampling Protocols

To produce the energy funnels for the fast folding proteins, we ran reverse diffusion from a Gaussian noise distribution at *t* = 1 with 200 reverse steps. For each noise level with *t <* 0.5, we ran 100 Langevin steps with *dt* = .001 before taking the next reverse step to descend to the next energy level. We ran 400 such trajectories for each protein. To reduce computational cost, we used the auxiliary score output for simulation instead of the true conservative score. The Langevin samples from the lowest noise level of each trajectory were then rescored with *t* = .05 to produce the final funnels and TICA plots. For steps with *t >* 0.1 we used a base ProteinEBM model finetuned on 1M frames of MD data with only the first and layers unfrozen. For steps *t <* 0.1 we used a ProteinEBM-x model trained on 1M frames MD with only the first and last layers unfrozen. We also ran the Langevin steps with a correction factor *T*_*train*_*/T*_*sim*_ = 300*/*350 to approximate the fact that the fast folder reference simulations were run at 350*K*, while our model was finetuned on simulations run at 300*K*.

For the Rosetta data set, we used a similar annealing protocol but with additional “refinement” stages to increase sample quality. For the initial sampling stage we followed the same protocol as for the fast folders, except using a model that was finetuned on 3M frames of MD with all parameters unfrozen. For refinement, we first filtered the best 5% of samples based on model energy, and resampled those structures with replacement with Boltzmann-weighted probabilities to generate 800 starting samples. Those samples were then renoised to *t* = 0.1, and denoised by Langevin annealing with a ProteinEBM-x model that was not finetuned on any MD data. During this annealing phase there were 20 reverse steps, with 10 Langevin steps at each noise level. During refinement we didn’t use a temperature scaling factor, and we used the conservative score from the model energy, which we found to produce more accurate structures. The refinement protocol was repeated three times, and the Langevin samples from the lowest noise level from all three refinement stages were pooled to create the final set of samples.

For the relaxation of native structures we noised them to *t* = .025 and then performed Langevin annealing using ProteinEBM-x with 5 reverse diffusion steps and 5 Langevin steps per noise level.

For direct folding simulations, we initialized the structure as a Ramachandran-random backbone. We simulated 10000 Langevin steps for Protein G and NuG2 and 25000 Langevin steps for Protein L, all with *dt* = .001. For Protein L, we also applied a small auxiliary potential of the form 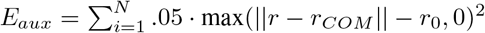, which pulls distant residues toward the protein’s center of mass when they exceed a baseline distance *r*_0_=10.75 Å. The auxiliary potential encourages the initially extended chain to quickly condense and fold.

## C Ablation of Energy for Ranking

As shown in our results, training an energy-based diffusion model results in an effective method to rank protein structures. A simpler alternative is to train an direct-score model and use the norm of the score as a pseudo-energy, which is related to the true energy:

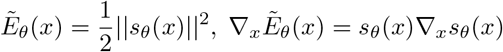

We implemented this method using the auxiliary direct score from our ProteinEBM models, and compared with the true energy. As seen in Figure S1, this method performed significantly worse.

**Figure S1:**
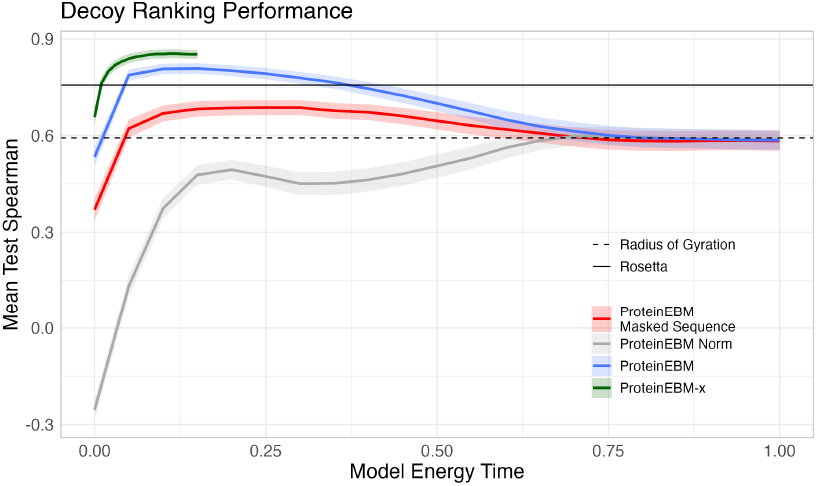
Rosetta test set spearman vs model energy time, including taking the norm of the score as a pseudo-energy.

## D Conformational Sampling Continued

As described in the main text, ProteinEBM accurately samples the native structures of fast-folding proteins and ranks the native-like conformations with the lowest energies. For NuG2, there is a second minimum at a strand-swapped version of the native structure which appears as the lowest-energy basin in some Langevin annealing runs (Figure S2). This result highlights the importance of both sufficient sampling and accurate energy ranking when using ProteinEBM for structure prediction.

**Figure S2:**
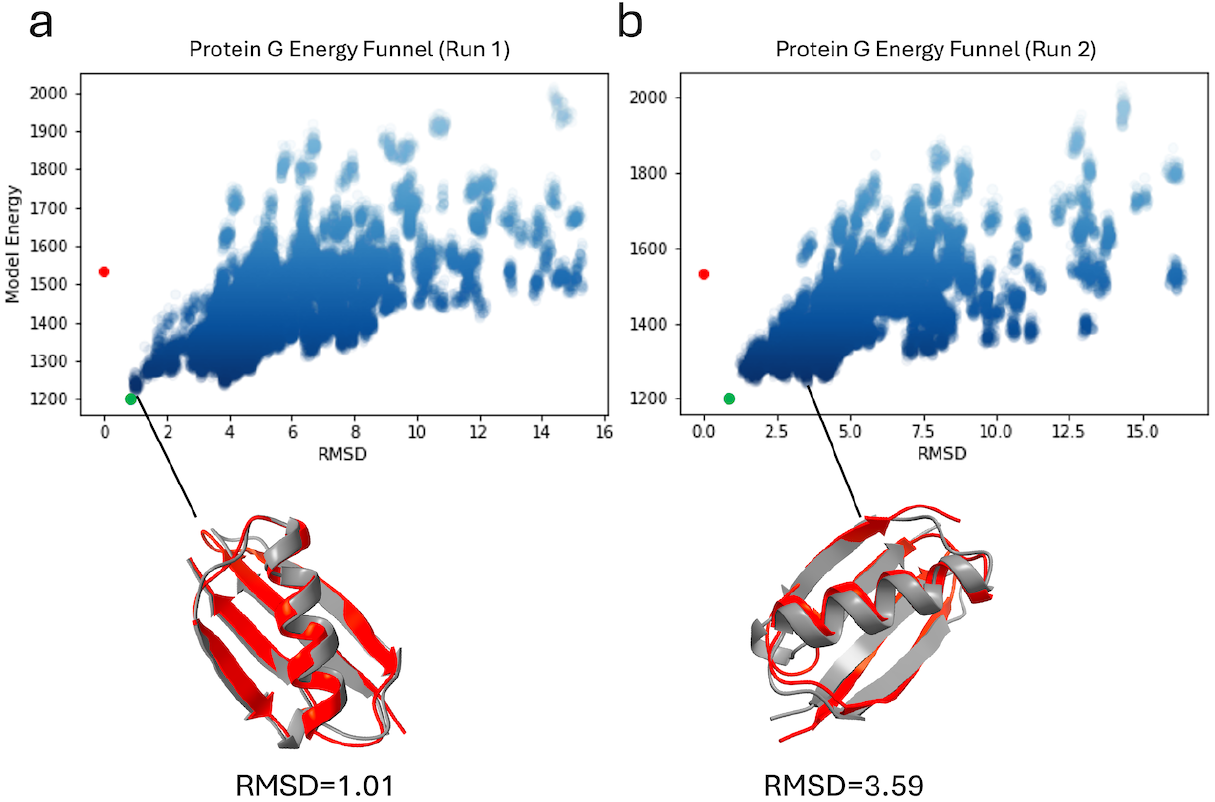
Two independent Langevin annealing runs, each with 400 trajectories. In a.) the lowest-energy sample adopts the correct fold and is extremely close to the native structure, while in b.) the lowest-energy sample has a swapped pair of strands compared to the native.

For Protein G and NuG2, the lowest energy sample from the direct folding simulations adopted the native fold (Figure S3a-b), while for Protein L the lowest-energy structure was a strand-swapped version of the native (Figure S3c). Despite this, some Protein L trajectories did reach the native structure, and we based our folding pathway analysis on those trajectories. The number of trajectories for each protein that reached the native structure (defined as RMSD <3 Å with correct secondary structure pairings) were 22/1200 for Protein G, 34/800 for NuG2, and 25/4800 for Protein L, which is consistent with the relative experimental folding speeds of these proteins.

**Figure S3:**
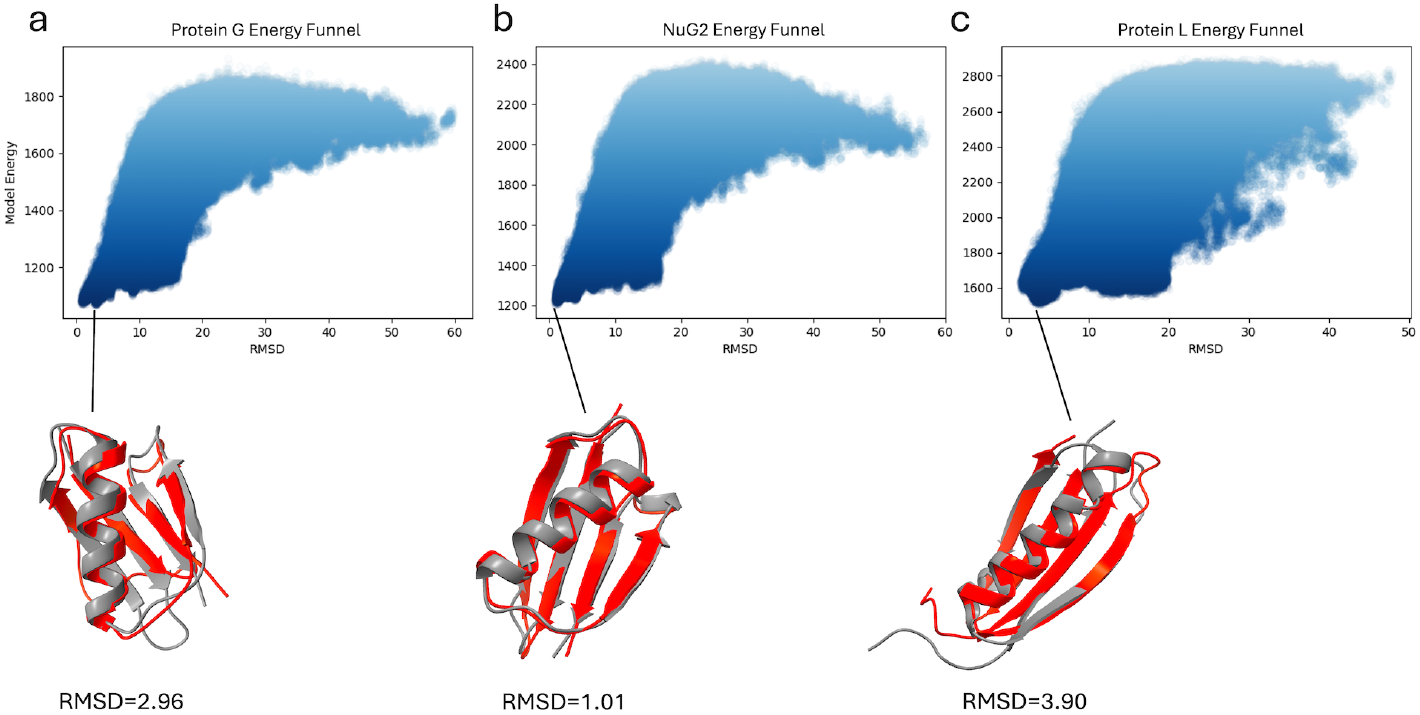
Energy funnels for direct folding simulations for a.) Protein G, b.) NuG2, and c.) Protein L. The lowest-energy samples for Protein G and NuG2 both have the same fold as the native, while for Protein L the lowest-energy structure has a swapped pair of strands.

## E Absolute Stability Prediction

In addition to predicting ΔΔ*G* of folding for mutated proteins, ProteinEBM-x can also be used in principle to predict absolute Δ*G* of folding for different proteins. To do this we computed

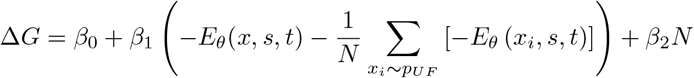

The approximation unfolded state energy uses the same estimator as in relative stability estimation. *β*_0_ and *β*_1_ are linear coefficients fit on top of the estimator to account for differences in units, *N* is the number of residues, and *β*_2_ models the per-residue loss in conformational entropy. We evaluated ProteinEBM on this task using data from Tsuboyama et al., following the analysis of Cagiada et al. [41, 49].

ProteinEBM-x showed mediocre performance at this task, with a Spearman correlation of .471 and a MAE of 1.06 kcal/mol. This is significantly worse than the results from ESM-IF in Cagiada et al. (*ρ* = .672, MAE = .96), although it is somewhat better than the naïve baseline of predicting based on the number of amino acids (*ρ* = .415, MAE = 1.12). We hypothesize that accurately predicting absolute folding energies may require a better model of the unfolded ensemble. ProteinEBM-x could likely be improved on this task by training directly on experimental stability measurements, as was done for BioEmu [23].

## F Designability Filtering

Machine learning methods for *de novo* protein design have rapidly proliferated in recent years. One common pipeline for structure-based design is to generate a candidate backbone, design a sequence for that backbone using ProteinMPNN or related models, and finally verify that the designed sequence is predicted to fold into the desired backbone [47]. In the final step, common computational thresholds for filtering designs include verifying that the predicted structure folds with *<* 2Å RMSD to the target backbone with high pLDDT or pTM.

**Figure S4:**
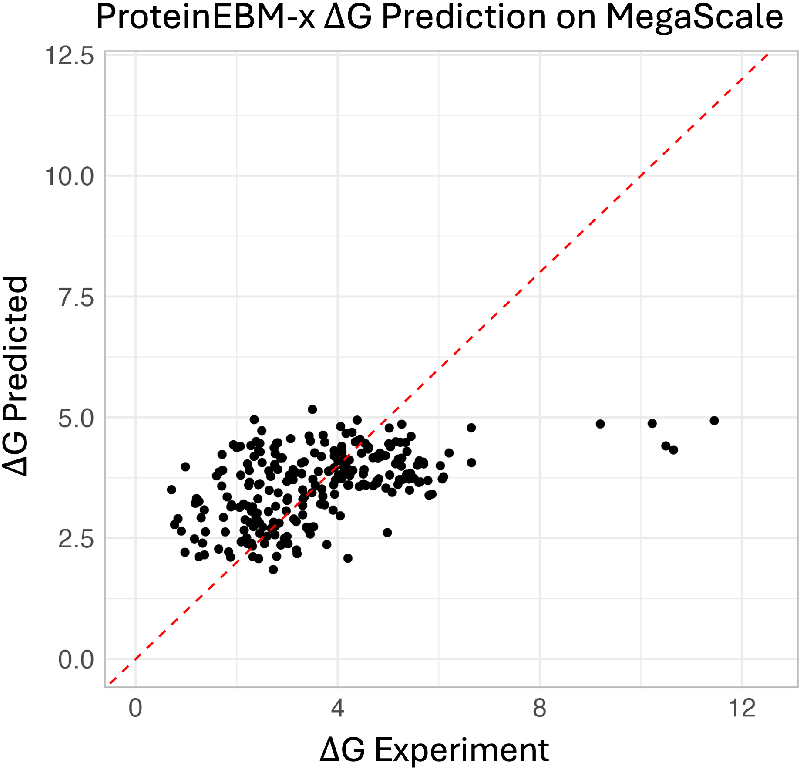
Evaluation of ProteinEBM-x Δ*G* prediction on MegaScale dataset.

One issue with this pipeline is that the refolding step can be very slow. In the recently published Boltzgen pipeline, refolding accounts for approximately 60% of the time spent generating designs [50]. To see if ProteinEBM-x could serve as a faster means of filtering designs, We generated 500 monomeric designs of length 200 using Boltzgen and evaluated the ProteinEBM-x energies of each designed backbone with its designed sequence. We found that ProteinEBM-x energies performed very well at detecting structures with high pTM (ROC AUC=0.952 for pTM > 0.7) and low refolding RMSD (ROC AUC=0.889 for RMSD < 2) as predicted by Boltz-2 (Figure S5). These results suggest that prefiltering with ProteinEBM could vastly speed up the refolding step of protein design pipelines like Boltzgen, as ProteinEBM-x is orders of magnitude faster than structure prediction methods like Boltz-2 (see Figure S7).

**Figure S5:**
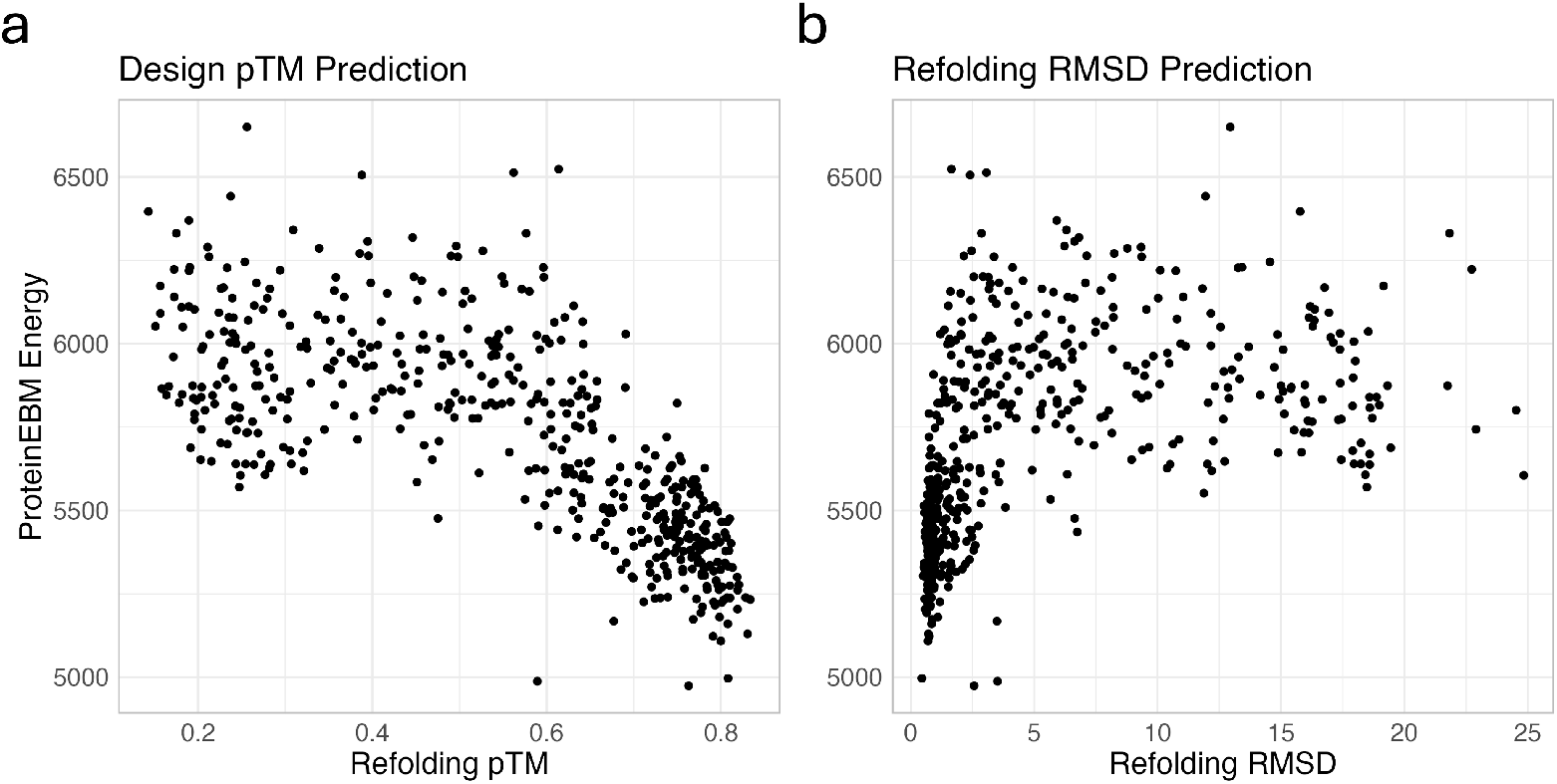
ProteinEBM-x can be used to filter designed proteins. a.) Correlation between ProteinEBM energy of design and Boltz-2 refolding pTM b.) Correlation between ProteinEBM energy of design and Boltz-2 refolding RMSD.

## G Conformational Biasing

One of the most interesting applications of EBMs is predicting the relative favorabilities of different conformational states. The free energy difference between two conformations (in units of *k*_*B*_*T*) can be approximated as − (*E*_*θ*_(*x*_1_, *s, t*) − *E*_*θ*_(*x*_2_, *s, t*)). As an initial exploration of this capability, we applied ProteinEBM-x to predicting the free energy difference between the open and closed states for various mutants of lipoic acid ligase (LplA). Cavanaugh et al. showed that ProteinMPNN can be used to predict which mutations bias LplA toward the open or closed state [51]. They computed the score log (*p*(*s*′ *x*_*open*_)*/p*(*s*′ *x*_*closed*_)) for various mutant sequences *s*′, with *p* computed using ProteinMPNN or another structure-to-sequence model. By Bayes’ rule, the following identity holds:

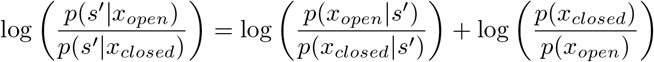

So the ProteinMPNN conformational biasing score is related by a constant offset to the true free energy difference log 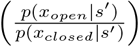. Therefore this score is mathematically correct for ranking purposes, but may not be accurate for predicting which conformation will dominate in absolute terms.

We conducted a similar analysis, ranking the mutations by −(*E*_*θ*_(*x*_*open*_, *s*′, *t*) −*E*_*θ*_(*x*_*closed*_, *s*′, *t*)), using *t* = 0.05 with the same model used for decoy ranking. Like the ProteinMPNN score, we found the ProteinEBM-x energy score to be positively correlated with experimental measurements of LplA promiscuity, which is correlated with time spent in the open state. Although ProteinEBM energies had significant predictive power, they were less correlated with promiscuous activity than the ProteinMPNN score.

**Figure S6:**
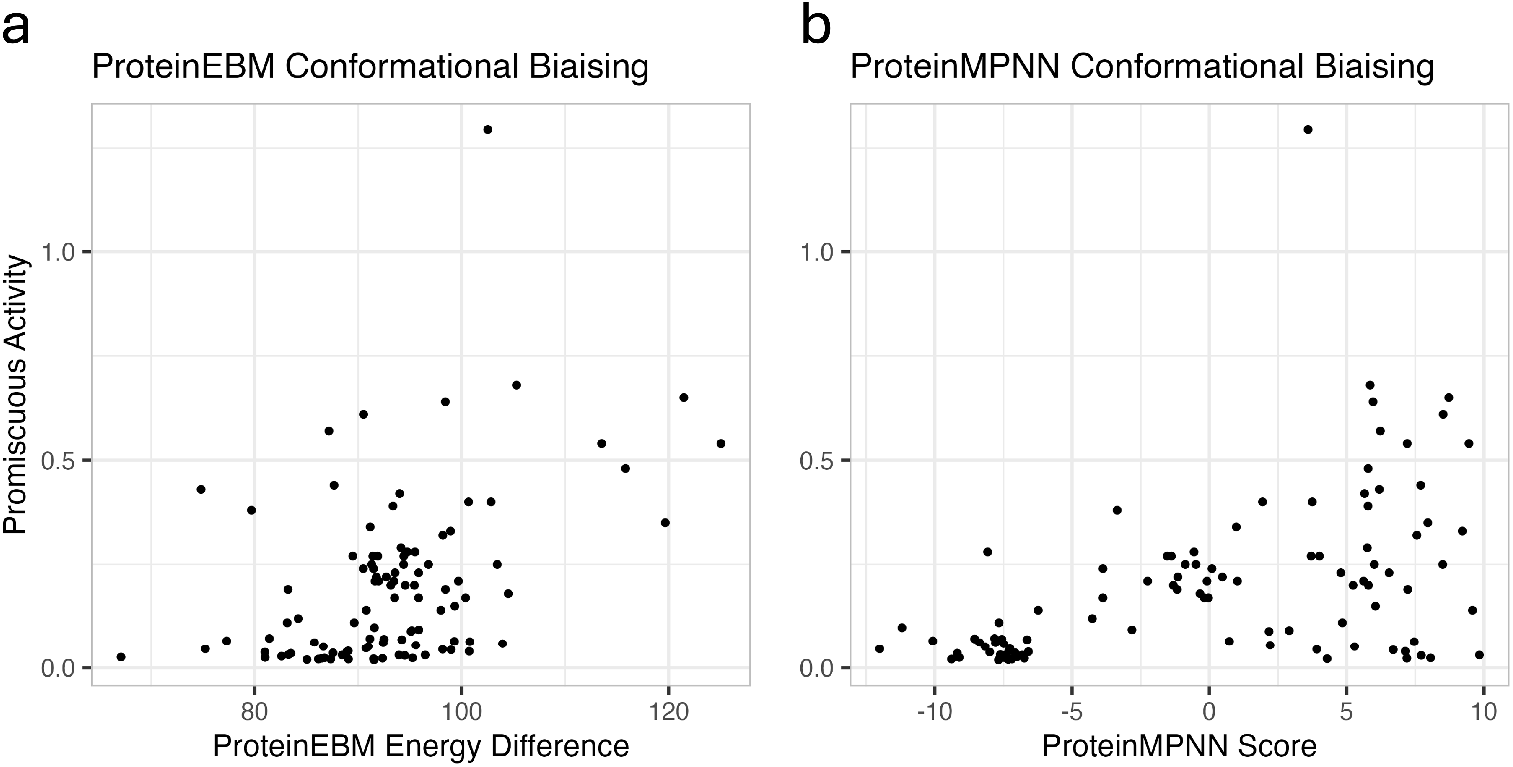
ProteinEBM-x can be applied to design conformationally biasing mutations. Correlation with LplA promiscuity for a.) ProteinEBM-x energy difference, b.) ProteinMPNN score.

In theory the ProteinEBM-x score should predict the free energy differences in absolute terms, unlike ProteinMPNN. However, the data from Cavanaugh et al. only reflect conformation occupancy via the measured promiscuity of LplA activity, which is correlated with time spent in the open state but does not reveal the absolute difference in state occupancies. Future analyses should explore whether ProteinEBM-x’s energy differences are accurate for proteins with known free energy differences between states.

## H Performance Analysis

To benchmark relative computational cost, we measured the runtimes of AlphaFold3, AlphaFold2, ProteinEBM, and Rosetta on proteins of various lengths (Figure S7). To avoid comparing Jax and Pytorch models, we used the OpenFold implementation for AlphaFold2, and used Boltz-1 as a surrogate for AlphaFold3. Based on these results, we calculated the number of samples from AlphaFold3 and AlphaFold2 needed to match the computational resources used for Langevin annealing with ProteinEBM. For ProteinEBM sampling we ran 400 initial annealing trajectories, each of which consisted of 200 reverse steps and 100 Langevin steps for each noise level under 0.5. This gives 400×(200+100×100) = 4, 080, 000 steps (See Appendix B.2 for details). We then conducted three refinement stages in which 800 trajectories were resampled and renoised to *t* = 0.1, and then denoised with 20 reverse steps and 10 Langevin steps per noise level for 3 × 800 ×(20 + 20 × 10) = 528, 000 steps. Since we used conservative scores for the refinement stages, we add an additional cost factor of 3 based on Figure S7, giving a total of 4, 080, 000 + 3 × 528, 000 = 5, 664, 000 total effective steps.

To compute conversion factors, we used the ratios of measured runtimes for a protein of size 128, which is representative of our test set. This relative runtime factor is 2 for the AF3 diffusion model, so with 200 diffusion steps per AF3 sample the number of computationally equivalent samples is 5, 664, 000*/*2*/*200 = 14, 160, which we rounded up to 15,000. To generate these samples, we ran AF3 with 150 seeds per target, generating 100 independent diffusion samples from each seed.

For AF2 the relative conversion factor is approximately 200 per model iteration. We used 3 recycles, so the equivalent number of samples is 5, 664, 000*/*200*/*4 = 7, 080, which we rounded up to 10,000. For AF2 we ran 2000 seeds per model across the 5 available AF2 monomer models with a pTM head.

**Figure S7:**
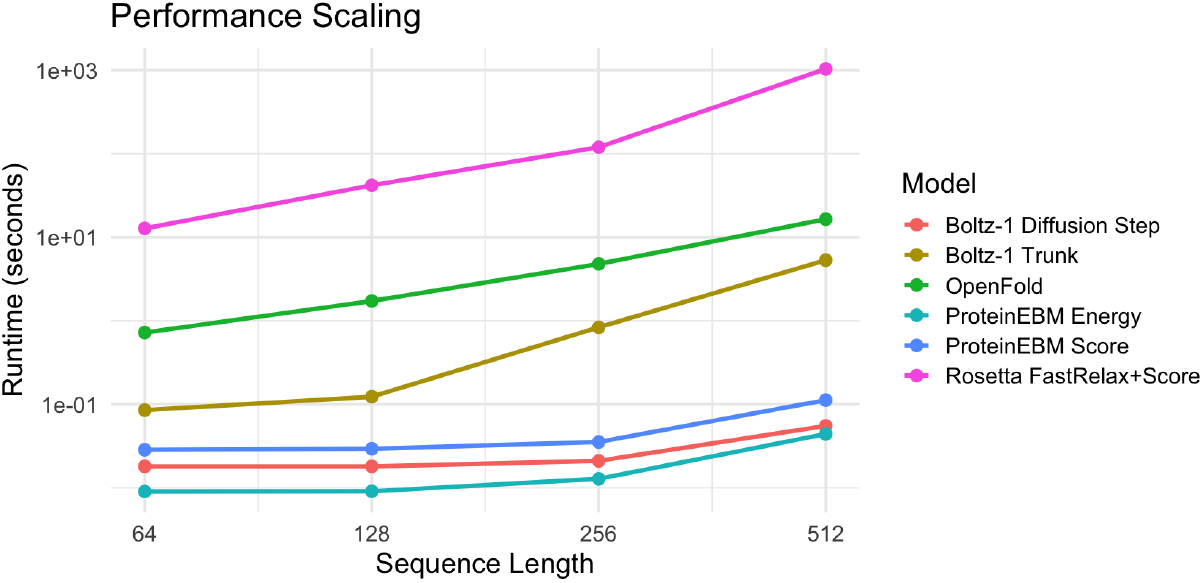
Runtime vs protein size for various protein modeling tools.

## I Effect of Sample Quantity on Prediction Accuracy

We chose to use 10,000 and 15,000 samples for our baseline comparisons against AF2 and AF3, respectively. Using bootstrapping, we estimated how top-1 accuracy increased as sample numbers were increased. The results of this analysis are shown in Figure S8. Interestingly, while the top-ranked AF2 TMScore increased significantly with increased sampling, this was not the case for AF3. However, the ground-truth highest TMScore (“Oracle” in Figure S8b) did increase steadily with increased sampling. This suggests that AF3’s confidence module may be less well calibrated than AF2, at least in the case of MSA-free monomer sampling.

**Figure S8:**
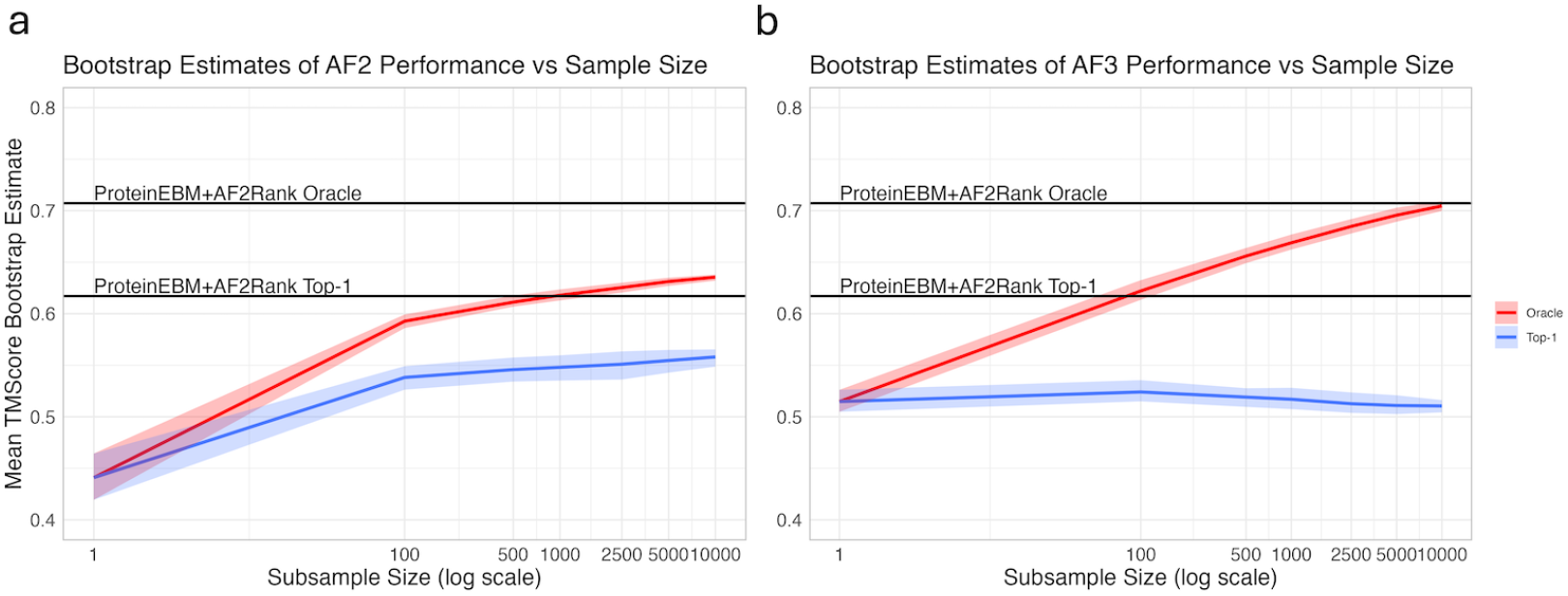
Examining the effect of sample size on AF2 and AF3 structure prediction. a.) Oracle TMScore and TMScore of top-ranked decoy for AF2 as number of samples is increased. b.) Oracle TMScore and TMScore of top-ranked decoy for AF3 as number of samples is increased. Ribbons are 95% bootstrap confidence intervals of the mean.

## J Failure Modes

During structure prediction, ProteinEBM exhibited a variety of failure modes on some targets. Two examples are shown in Figure S9.

**Figure S9:**
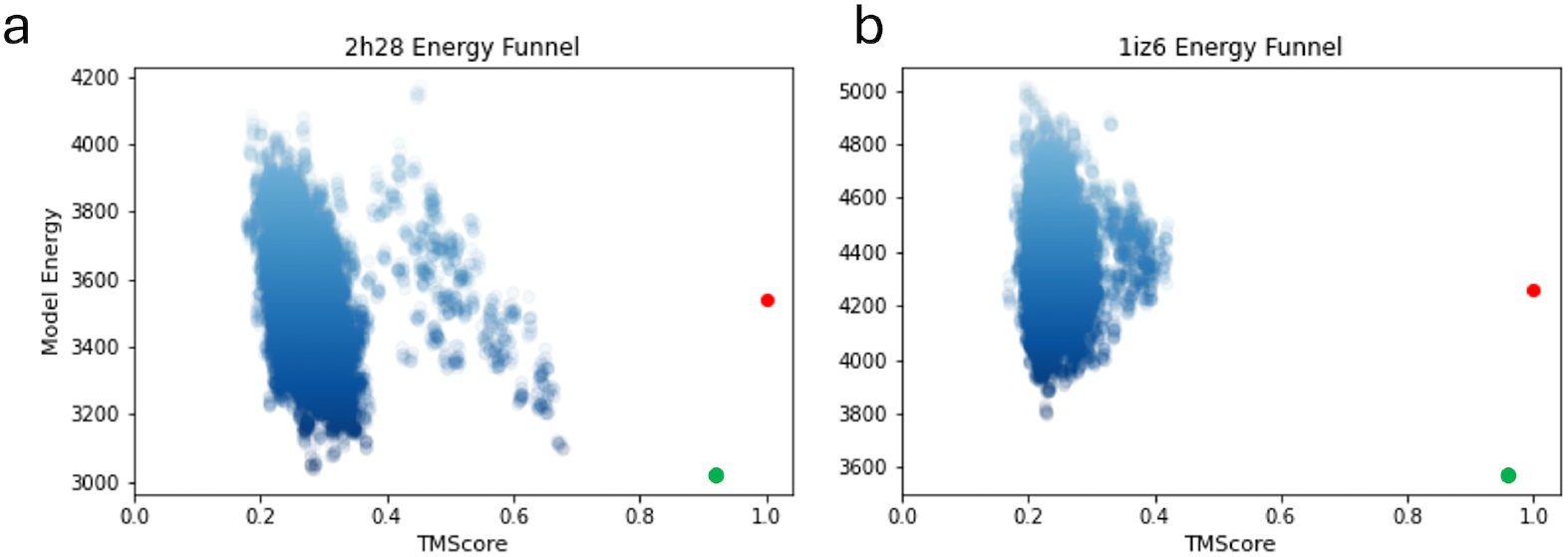
Energy funnels for failed prediction targets, with native structure as red dot and relaxed native structure as green dot. a.) Example where ProteinEBM explored a structure close to the native (TMScore ≈ 0.7), but erroneously scored a much less accurate structure with lower energy. b.) Example where the model failed to explore close to the native structure, although the native structure had a far lower energy than the structures that were actually sampled.

